# Tumor-agnostic transcriptome-based classifier identifies spatial infiltration patterns of CD8+ T cells in the tumor microenvironment and predicts clinical outcome in early- and late-phase clinical trials

**DOI:** 10.1101/2023.10.02.560086

**Authors:** Andreas Roller, Iakov I Davydov, Petra C Schwalie, Martha L Serrano-Serrano, Astrid Heller, Nicolas Staedler, Cláudia S Ferreira, Gabriele Dietmann, Irina Klaman, Alberto Valdeolivas, Konstanty Korski, Michael A Cannarile

## Abstract

**Background:** The immune status of a patient’s tumor microenvironment (TME) may guide therapeutic interventions with cancer immunotherapy and help identify potential resistance mechanisms. Currently, patients’ immune status is mostly classified based on CD8+ tumor-infiltrating lymphocytes. An unmet need exists for comparable and reliable precision immunophenotyping tools that would facilitate clinical treatment-relevant decision-making and the understanding of how to overcome resistance mechanisms.

**Methods:** We systematically analyzed the CD8 immunophenotype of 2023 patients from 14 phase I–III clinical trials using immunohistochemistry (IHC) and additionally profiled gene expression by RNA-sequencing (RNA-seq). CD8 immunophenotypes were classified by pathologists into CD8-desert, CD8-excluded or CD8-inflamed tumors using CD8 IHC staining in epithelial and stromal areas of the tumor. Using regularized logistic regression, we developed an RNA-seq-based classifier as a surrogate to the IHC-based spatial classification of CD8+ tumor-infiltrating lymphocytes in the TME.

**Results:** The CD8 immunophenotype and associated gene expression patterns varied across indications as well as across primary and metastatic lesions. Melanoma and kidney cancers were among the strongest inflamed indications, while CD8-desert phenotypes were most abundant in liver metastases across all tumor types. A good correspondence between the transcriptome and the IHC-based evaluation enabled us to develop a 92-gene classifier that accurately predicted the IHC-based CD8 immunophenotype in primary and metastatic samples (area under the curve (AUC) inflamed = 0.846; excluded = 0.712; desert = 0.855). The newly developed classifier was prognostic in The Cancer Genome Atlas (TCGA) data and predictive in lung cancer: patients with predicted CD8-inflamed tumors showed prolonged overall survival (OS) versus patients with CD8-desert tumors (hazard ratio [HR] 0.88; 95% confidence interval [CI]: 0.80–0.97) across TCGA, and longer OS upon immune checkpoint inhibitor administration (phase III OAK study) in non-small-cell lung cancer (HR 0.75; 95% CI: 0.58–0.97).

**Conclusions:** We provide a new precision immunophenotyping tool based on gene expression that reflects the spatial infiltration patterns of CD8+ lymphocytes in tumors. The classifier enables multiplex analyses and is easy to apply for retrospective, reverse translation approaches as well as for prospective patient enrichment to optimize the response to cancer immunotherapy.

**HIGHLIGHTS:** *What is already known on this topic:* T-cell infiltration, most commonly classified based on CD8+ T cell immunohistochemistry (IHC) staining, and various tumor microenvironment (TME)-specific resistance mechanisms, can impact response rates to cancer immunotherapy.

*What this study adds:* Our data provide new insights into the impact of tumor excision location and indication on the immune composition of the TME. We developed a transcriptome-based classifier that could accurately predict different spatial CD8+ T-cell infiltration patterns in the TME. We demonstrate the prognostic and predictive value of the classifier across independent patient cohorts (phase I to phase III trials).

*How this study might affect research, practice or policy:* Our new RNA-based tool provides a surrogate read-out for spatial IHC-based CD8 infiltration patterns, is easy to use and broadly applicable for both retrospective and prospective patient enrichment to enhance the effectiveness of cancer immunotherapy.

## INTRODUCTION

Cancer immunotherapy (CIT) has been widely integrated into routine clinical practice for many tumor indications, helping to move from a disease-centric approach to a more personalized care with substantially improved outcomes.^1 2^ However, varied response and resistance rates can be attributed to several factors, including paucity of T-cell infiltration in the tumor microenvironment (TME) and tumor-intrinsic resistance mechanisms.^3–5^ Improved understanding of resistance mechanisms, along with reliable prognostic and predictive biomarkers, would help achieve the full potential of CIT.^6 7^ Reliable predictive biomarkers could eliminate administration of immunotherapy to unsuitable patients, minimizing the associated risk of toxicity, and reducing costs.^8 9^

Many biomarkers have been investigated, with various limitations, challenging their implementation into routine clinical practice.^1 7 10^ Among these, the immune status of a patient is commonly assessed by analyzing the TME at baseline using immunohistochemistry (IHC) and classifying the spatial infiltration of CD8+ T cells in tumors as ‘CD8-inflamed’ (a high rate of infiltration, typically showing significant immune cell diversity), ‘CD8-excluded’ (T cells are retained in the stroma) or ‘CD8-desert’ (very limited infiltration in intra-tumoral stroma and tumor areas).^2 11 12^ A significant correlation between CD8-inflamed tumors and improved clinical response to CIT has been reported^2 13 14^, including for pembrolizumab treatment of metastatic melanoma^15^ and oral squamous cell carcinoma.^16^ IHC is useful to broadly categorize tumors, but risks simplifying the complex genetic and immune picture inside the TME.^6^ For example, with lymphocytes, IHC often only reports CD8 positivity and may not identify other pathological pathways (e.g. altered tumor metabolic pathways); more complex biomarker approaches are thus needed to further identify the dynamics of tumorigenesis. IHC also assumes all centers have equal access to reliable assays, appropriately curated samples, and equivalent means of interpreting results.^6^ Therefore, it may not always be the optimal approach to delineate tumor immunophenotypes and predict therapeutic response, leaving an unmet need for reliable and cohesive biomarkers to help guide patient care.

Gene expression-based models have been developed to classify tumor immunophenotypes, for example, by using RNA sequencing (RNA-seq) to analyze gene expression related to CD8+ T-cell activity. In line with the IHC-based assessments, the T-effector gene signature was also associated with an improved efficacy of atezolizumab versus docetaxel in patients with non-small-cell lung cancer (NSCLC).^17–19^ Similar signatures have been used to generate risk scores correlated with CD8+ T-cell infiltration^13^, which could predict immunotherapy efficacy, potentially enabling early identification of non-responders.^20^ Importantly, gene expression profiling by RNA-seq can generate comprehensive and cost-effective datasets, with demonstrated concordance between IHC, quantitative real-time polymerase chain reaction (RT-PCR) and gene expression microarrays.^21^ An easy-to-implement gene expression-based classifier reflecting the spatial infiltration of CD8+ T cells in the TME is needed to support large-scale analyses of clinical datasets, enabling retrospective and reverse translation analyses across clinical studies. However, previous attempts at identifying RNA-seq signatures of immunophenotypes and their predictive and prognostic values were limited to specific cancer types.^20 22^

Here, we describe the development of a transcriptome-based classifier that accurately identifies spatial infiltration patterns of CD8+ T cells in the TME across indications and excision locations. This classifier is able to predict clinical outcomes in early- and late-phase clinical trials, which has significant implications for the use of CIT.

## METHODS

### Patient samples

Formalin-fixed and paraffin-embedded tumor tissue collected from 11 unpublished phase I/II clinical trials (**supplemental table S1**) before treatment start were retrospectively analyzed. Tumor metastases in the liver and lymph nodes were measurable and assessable as target lesions, allowing an immune score to be calculated.^23^

Additionally, IHC-based CD8 immunophenotypes were scored for samples extracted from three previously published open-label, phase II–III trials (NCT02008227; NCT02108652; NCT02302807), from which RNA-seq data were also available.^24–26^

All clinical studies were conducted in accordance with the principles of the Declaration of Helsinki and Good Clinical Practice Guidelines. Written informed consent was collected from all enrolled patients.

Data from The Cancer Genome Atlas (TCGA) Program (National Cancer Institute; available at https://portal.gdc.cancer.gov/) were used to test the gene expression-based classifier on a large dataset, processed via the recount3 resource.^27^

### CD8 immunophenotypes: IHC classification

CD8/KI67 slides were cut (2.5µm thickness) and stained in-house (Roche Innovation Center Munich, Penzberg, Germany). Slides were scanned at 20x using the Ventana iScan HT and the resulting whole-slide images were sent to CellCarta (formerly known as HistoGeneX [Antwerp, Belgium]) for immunophenotyping assessment/visual scoring. Phenotype scoring (or density proportion scoring, adapted from Galon and Lanzi 2020^28^) was based on CD8/KI67 IHC levels in tumor epithelial and stroma areas by CellCarta, although only CD8 levels were analyzed in our study. Based on a combined density score in the intra-epithelial (IE) and intra-tumoral stroma (ITS) score compartments, patient samples were classified as (**figure 1A**): (i) CD8-inflamed, with CD8+ cell counts > 500 cells/mm^2^ in the intraepithelial compartment with IE2 + IE3 ≥ 20%; (ii) CD8-excluded, with ≤ 500 cells/mm^2^ in the intraepithelial and > 50 cells/mm^2^ in the stromal compartment ITS0 + ITS1 ≥ 80% and ITS2 + ITS3 < 20%; (iii) CD8-desert, with ≤ 50 cells/mm^2^ in the stromal compartment IE0 + IE1 ≥ 80% & IE2 + IE3 < 20% and ITS0 + ITS1 < 80% and ITS2 + ITS3 ≥ 20%.

**Figure 1.**
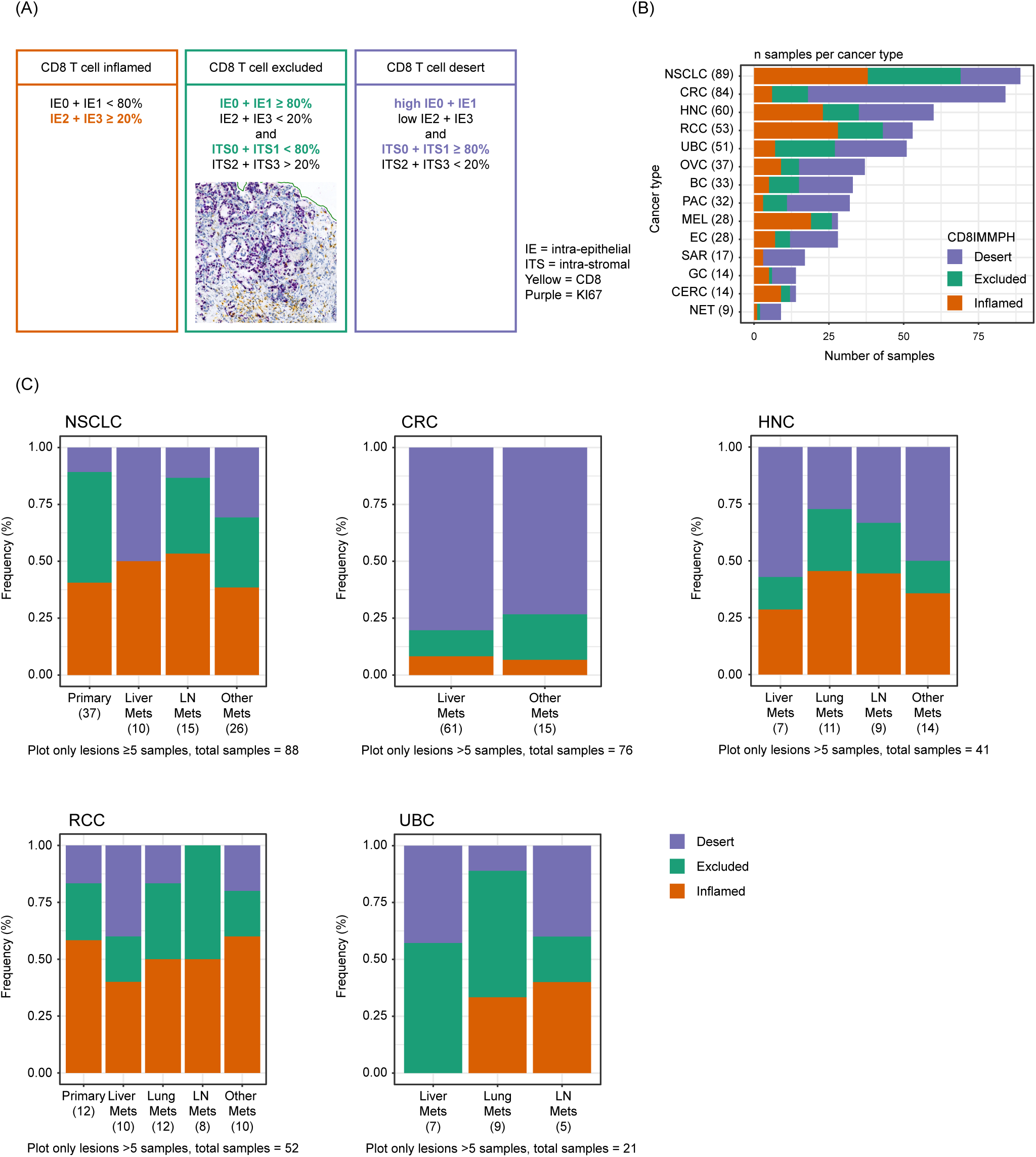
Definition and prevalence of CD8 immunophenotype categories. (A) CD8-inflamed was defined as CD8+ cell counts > 500 cells/mm^2^ in the intraepithelial compartment with IE2 + IE3 ≥ 20%. CD8-excluded was defined as ≤ 500 cells/mm^2^ in the intraepithelial and > 50 cells/mm^2^ in the stromal compartment ITS0 + ITS1 ≥ 80% and ITS2 + ITS3 < 20%. CD8-desert was defined as ≤ 50 cells/mm^2^ in the stromal compartment IE0 + IE1 ≥ 80% & IE2 + IE3 < 20% and ITS0 + ITS1 < 80% and ITS2 + ITS3 ≥ 20%. Prevalence of immunophenotype by (B) indication and (C) tumor excision location. The “Other mets” category refers to excision sites of minor frequency (each n ≤ 4): abdominal cavity, sinus, adrenal gland, pleura, skin and soft tissue. Tumor types with n < 5 samples (25 different categories, e.g., bone, fallopian tube, pelvis, thymus, testis; most of them n=1) are not included. IE, intraepithelial; ITS, intrastromal; LN, lymph node; mets, metastases; NSCLC, non-small-cell lung cancer; UBC, urinary bladder cancer.

### RNA-seq data processing

All samples were further analyzed for genome-wide RNA expression using RNA-seq, performed on macro-dissected tissue of the tumor area (i.e. normal tissue was excluded). RNA-seq data were analyzed using HTSeqGenie^29^ in BioConductor^30^ as follows: first, reads with low nucleotide qualities (70% of bases with quality < 23) or matches to rRNA and adapter sequences were removed. The remaining reads were aligned to the human reference genome GRCh38.p10 using GSNAP^31^ version ‘2013-10-10-v2’, allowing maximum two mismatches per 75 base sequence (parameters: ‘-M 2 -n 10 -B 2 -i 1 -N 1 -w 200000 -E 1 --pairmax-rna=200000 --clip-overlap’). Transcript annotation was based on the Gencode database (human: GENCODE 27). To quantify gene expression levels, the number of reads mapping unambiguously to the exons of each gene was calculated. RNA-seq values were transformed using variance stabilizing transformation and DESeq2 version 1.6.3 (Bioconductor, Massachusetts, USA). Lowly expressed genes and those with vastly different expression levels between TCGA and the rest of the data were excluded to allow the classifier to be applied to TCGA. Batch correction was subsequently performed: principal component analysis (PCA) transformation was used on a selected subset of lung cancer samples to identify components corresponding to the batch-effect (i.e., difference in mean and variance). Finally, the effect of those components on individual genes was identified and an inverse transformation was applied to the whole TCGA dataset.

Signature scores corresponding to rank-biserial correlation values were produced using BioQC version 3.15.^32^ Cell-type-specific signatures were previously derived from single-cell RNA-seq data,^33^ while pathway-specific signatures were available from the MsigDB “Hallmark” collection (Molecular Signature Database [MSigDB], release 7.4^34^). Pairwise differential expression analyses were performed to characterize gene expression differences between the three phenotypes. A linear model was fitted using the limma-voom approach defined by Law et al.^35^ with a threshold for log fold change of 1.5 and corrected false discovery rate (FDR) of 0.05. Biopsy location (liver, kidney, lung, urothelial, lymph node) was added as a covariate in the model. Gene set enrichment was calculated using a competitive gene set test (*limma::camera*) with FDR < 0.05. Differential expression results are provided as **supplemental table S2** and gene set enrichment results in **supplemental table S3**.

### Classifier development

To ensure adequate classifier performance estimates, the data (N=2023: n=628 from phase I/II trials and n=1395 from the three additional phase II/III trials) were split into three portions: training (71%; n=1438), validation (8%; n=158), and test (21%; n=427). To build the classifier, lasso regression was implemented in the glmnet R package^36^ via mlr3^37^ was used. Nested cross-validation was used to identify the optimal value of lambda. Variance-stabilization transformed gene expression values were selected as an input for the classifier. TCGA RNA-seq samples were batch-corrected to match the training dataset. To explore the most important features for classification, a set of classifiers were trained that used: (i) all genes (GENCODE 27, excluding genes that were filtered out, see section RNA-seq data processing); (ii) only immune-expressed genes, as derived from single-cell RNA-seq expression datasets available in besca;^33 38–40^ (iii) cell type-specific signatures scores,^33^ (iv) hallmark pathway signature scores (MSigDB, release 7.4^34^); (v) or cancer-specific signature scores (C4 and C6, MSigDB, release 7.4^34^). A 10-fold cross-validation was repeated 100 times (on random data splits) to measure the feature importance of individual genes. For every classifier parameter, the number of times it was present in one of the models and its coefficient value were recorded. The test fold was used to assess the classifier performance. Samples from the phase III OAK study (NCT02008227; n=352) were only used in the test fold to control for classifier generalizability and translatability. Using a study for the test fold that was not included in training ensures that the prediction performance is a good approximation of classifier generalizability to completely new, unseen data. Inter-gene correlation between gene signatures was also calculated (see **supplemental figure S1**).

### Single-cell RNA-seq and spatial transcriptomics expression analysis

To explore which cell types express the top genes retained in the final classifier, previously published colorectal cancer (CRC),^40^ lung cancer^41^ and liver cancer^42^ datasets were reprocessed with the besca standard workflow and cell annotation workflow.^33^ To explore the spatial distributions of individual genes, publicly available 10x Visium data from CRC^43^ were reprocessed as described previously.^44^

### Statistical analyses

The R package limma was used for the differential gene expression analysis.^35^ The RTCGA package version 1.26.0 was used to extract clinical information from TCGA.^45^ The survminer R package version 0.4.9 package was used to fit Cox proportional hazard models.^46^ The meta version 6.1-0 package was used to produce forest plots.

## RESULTS

### CD8 immunophenotypes vary across indications and tumor excision locations

Tumor samples from 628 patients enrolled into 11 phase I/II clinical trials were systematically analyzed by IHC and RNA-seq (see **supplemental table S1** for a detailed overview). Patient baseline characteristics are reported in **supplemental table S4**: median age was 60 years, and 43% of patients were female. The most common diagnoses were NSCLC (n=89), CRC (n=84), and head and neck carcinoma (HNC) (n=60). A total of 79% of patients were treatment naïve.

Samples were classified based on CD8 IHC levels in tumor epithelial and stroma areas as CD8-inflamed (31%), CD8-excluded (23%), and CD8-desert (46%) (**figure 1A**). The distribution of immunophenotypes was highly variable across indications and tumor excision locations (**figures 1B–C**). As expected from previous reports, the majority of NSCLC, HNC and urinary bladder carcinoma (UBC) samples were CD8-inflamed and/or CD8-excluded. There was also a high prevalence of CD8-inflamed phenotypes in melanoma (MEL) and renal cell carcinoma (RCC) samples, whereas CD8-desert phenotypes were predominant in CRC. There was a higher relative fraction of CD8-desert phenotypes in liver metastases across all tumor types: approximately 50% (versus less than 25% in lymph node metastases or in the primary tumor) of liver metastases from NSCLCs had a CD8-desert phenotype. Similarly, the proportion of CD8-desert phenotype in liver metastases was 40% (vs 10% in lung metastases) for UBCs, and 80% (vs 60% in the primary tumor) for CRCs. Trends were also consistent for RCC and HNC (**figures 1B–C**).

### CD8 immunophenotypes show distinct transcriptional profiles

The transcriptional characteristics of the three CD8 immunophenotypes were first explored with a focus on cell type and pathway-specific gene signatures previously positively or negatively associated with immune infiltration and/or response to cancer immunotherapy (**figure 2A**). Globally, gene expression was similar, albeit with magnitude differences, in the CD8-inflamed and CD8-excluded groups but showed rather distinct patterns from CD8-desert samples, across indications and excision locations (**figure 2A**). Gene expression related to CD8+ T cells, cytotoxicity, exhaustion, as well as interferon (IFN) response, was highly enriched in CD8-inflamed samples, intermediate in CD8-excluded, and low in CD8-deserts, in line with the abundance and activity of CD8+ T cells in CD8-inflamed samples (**figures 2A–B; supplemental figure S2**). In contrast, expression levels related to immune populations previously associated with repression of T-effector activity, such as regulatory T cells, macrophages, or myeloid cells, showed more similar expression in inflamed and excluded samples, and markedly lower expression in CD8-desert samples. Stromal-specific genes, including fibroblast and endothelial markers, as well as angiogenic and transforming growth factor-β (TGF-β) pathways, showed excluded-enriched expression (**figure 2B; supplemental figure S2**).

**Figure 2.**
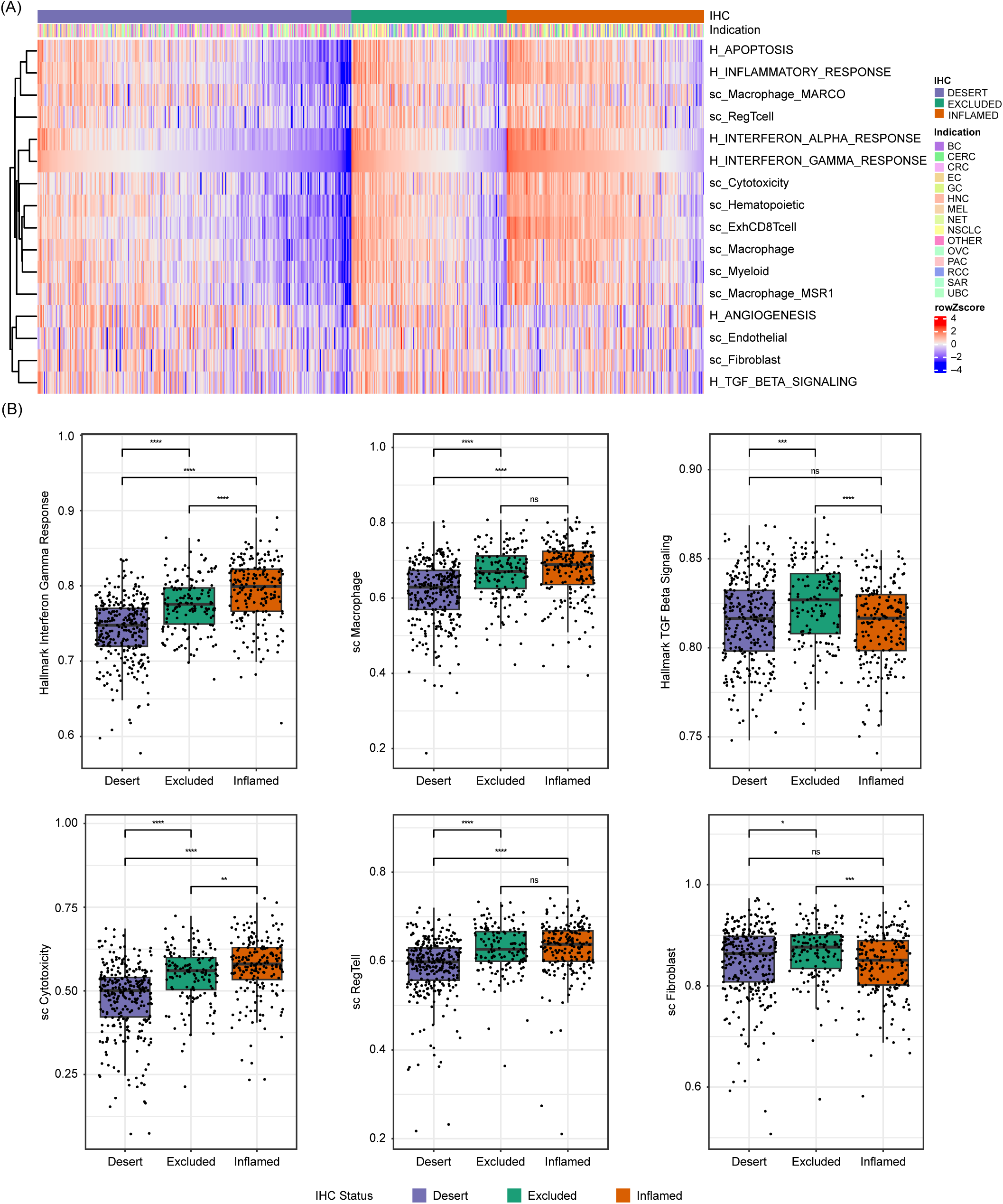
Inflamed, excluded and desert samples show distinct transcriptional profiles. (A) Heatmap showing relative signature scores per sample, sorted according to CD8 immunophenotype and “Interferon gamma” signature scores; cancer types are displayed. (B) Selected signatures scores across CD8 immunophenotype classes. “H_” represents pathway-specific signatures (Hallmark), while “sc_” represents cell-type specific signatures. IHC, immunohistochemistry; ns, not significant.

To systematically characterize gene expression differences between CD8 immunophenotypes, we next performed pairwise differential expression analyses. Given the observed heterogeneity of our phase I/II cohort, we included an additional set of 1395 samples (2023 samples in total) from three phase II–III trials to better control for the observed indication and excision location-induced variability. The greatest differences were between CD8-inflamed and CD8-desert samples, consistent with the signature-based analysis (**figure 3**; **supplemental table S3**). Pairwise differential expression analyses also showed that T-effector and IFN-ɣ pathway-related gene expression, including *CXCL9, CXCL10, IFN-ɣ, CCL5, ITGAE, LAG3, FASLG,* and *TAP1*, was most highly enriched in CD8-inflamed samples (**figure 3B–D; supplemental figure S3A**). The same genes were significantly lower expressed in CD8-excluded versus CD8-inflamed samples, but more highly expressed between CD8-excluded and CD8-desert samples; this was consistent across indications.

**Figure 3.**
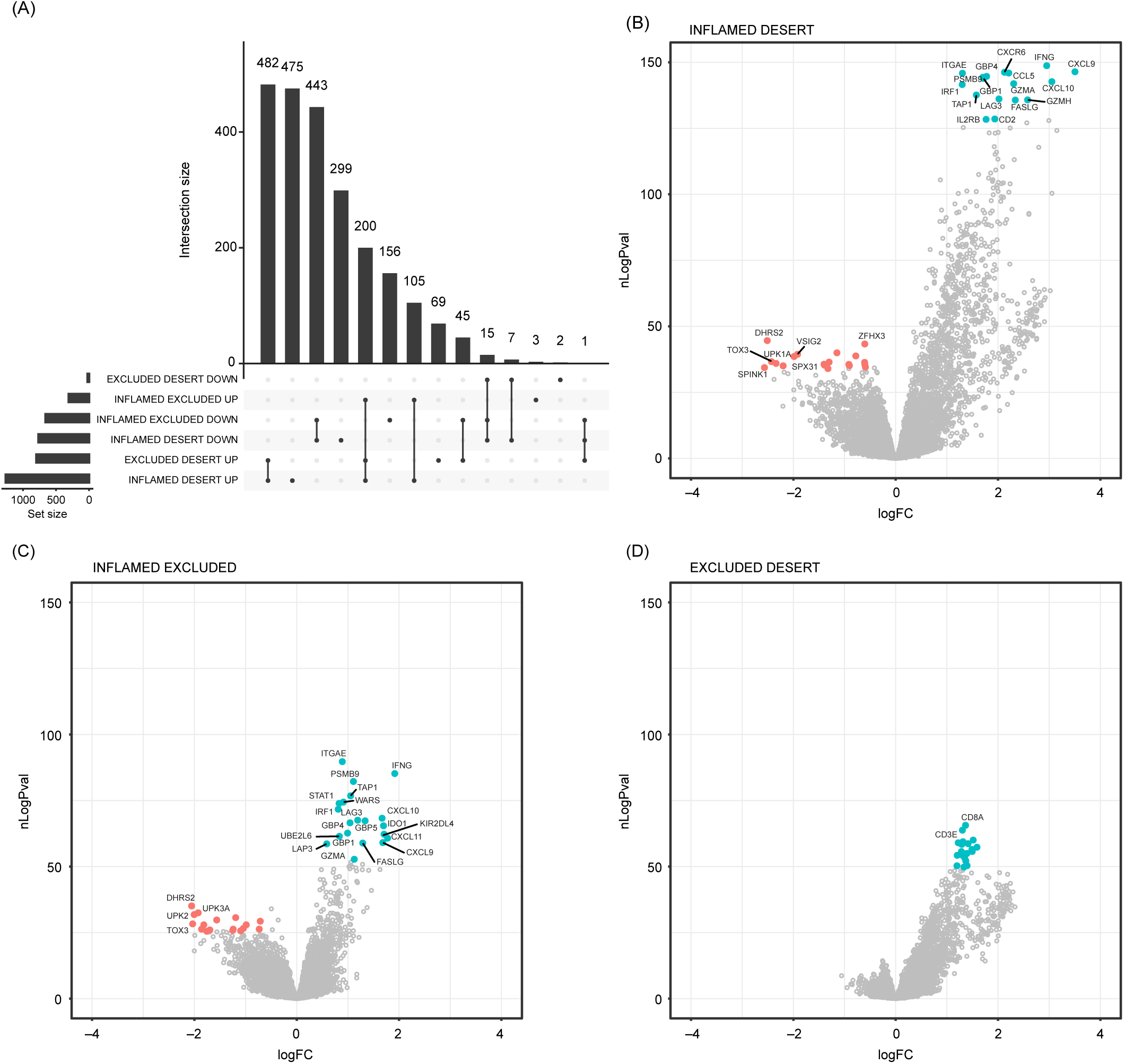
Systematic characterization of gene expression differences across CD8 immunophenotypes. (A) Overlaps between the genes significantly differentially expressed across all pairwise comparisons. (B–D) Volcano plots highlighting the most strongly differentially expressed genes in inflamed versus desert (B), inflamed versus excluded (C), and excluded versus desert (D) tumors. Upregulated genes are in blue, downregulated genes are in red. CD8IMMPH, CD8 immunophenotype; logFC, log fold change; nLogPval = −log10 (P value).

Few (n=22) genes were more highly expressed in CD8-desert samples than in both CD8-inflamed and CD8-excluded samples (desert-enriched): such genes were also often more highly expressed in CD8-excluded versus CD8-inflamed samples (**supplemental figure S3B**). Similarly, only 45 genes were more highly expressed in CD8-excluded samples versus both CD8-inflamed and CD8-desert samples. These included TGF-β signaling genes such as *PLN*, *C7*, *ADH1B*, *OGN*, *SCRG1*; the Wnt signaling genes *SFRP4*, *SFRP1*, *SFRP2*, and genes characteristic of smooth muscle (*CNN1*, *ACTC1*, *DES*) and mast cells (*CTSG*, *MS4A2*). Consistently, fibroblast, endothelial, and mast cell signatures were significantly higher enriched in CD8-excluded versus both CD8-inflamed and CD8-desert samples (**supplemental figure S3C; supplemental table S3**).

### IHC-based CD8 immunophenotypes can be accurately predicted from transcriptomic data

Given the large number of differentially expressed genes across the three immunophenotype classes and their high consistency across indications and excision locations, we hypothesized that the transcriptomic information may be sufficient to distinguish CD8-inflamed, CD8-excluded and CD8-desert samples, in the absence of IHC data. To test this hypothesis, as well as to better understand molecular mechanisms driving immunophenotypes by narrowing down the list of genes/pathways critical for their separation, we developed a set of classifiers using distinct input features. We used either: (i) all detected genes; (ii) genes typically only expressed in immune cells; (iii) a small set (97) of cell-type specific signatures scores previously derived from single-cell RNA-seq data;^33^ (iv) hallmark pathway signatures (MSigDB);^34^ or (v) cancer-specific signatures (C4 and C6, MSigDB)^34^ as input features, as described in the **methods**. We developed all classifiers using the full 2023 patient sample set from 14 phase I–III trials while performing area under the curve (AUC) evaluation on 5-fold cross-validation.

All classifiers demonstrated high performance in identifying CD8-inflamed and CD8-desert samples with a mean AUC > 0.8, while CD8-excluded samples were more challenging to delineate (AUC 0.70–0.76; **figure 4A**). The best performance was obtained when using either all genes or only immune-related genes, consistent with a large fraction of the signal being derived from the immune compartment. The classifier trained on cell-type-specific signature scores showed comparable performance, while pathway signatures and cancer-specific gene expression-trained classifiers showed the lowest accuracy (**figure 4A**). T-effector-related signatures were retained in this classifier: high signature values associated positively with the CD8-inflamed phenotype (e.g., “Macrophage_CXCL9”, “ExhCD8Tcell”, “Interferon-alpha/gamma response”) or negatively with the CD8-desert phenotype (e.g., “T cell”, “Cytotoxicity”, “Allograft rejection”) (**supplemental figure S4A**). By contrast, stromal-related expression (”Fibroblast”), known immunosuppressive pathways (TGF-β-signaling), or pathways related to tumor survival and aggressiveness (NOTCH signaling) were negatively associated with the CD8-inflamed phenotype. Only a few signatures were discriminatory for the CD8-excluded phenotype, most notably “Regulatory T cell”, “Exhausted B cells”, “Adipocytes”, “Adipogenesis”, “Androgen response”, and “PI3K_AKT_MTOR_Signaling”.

**Figure 4.**
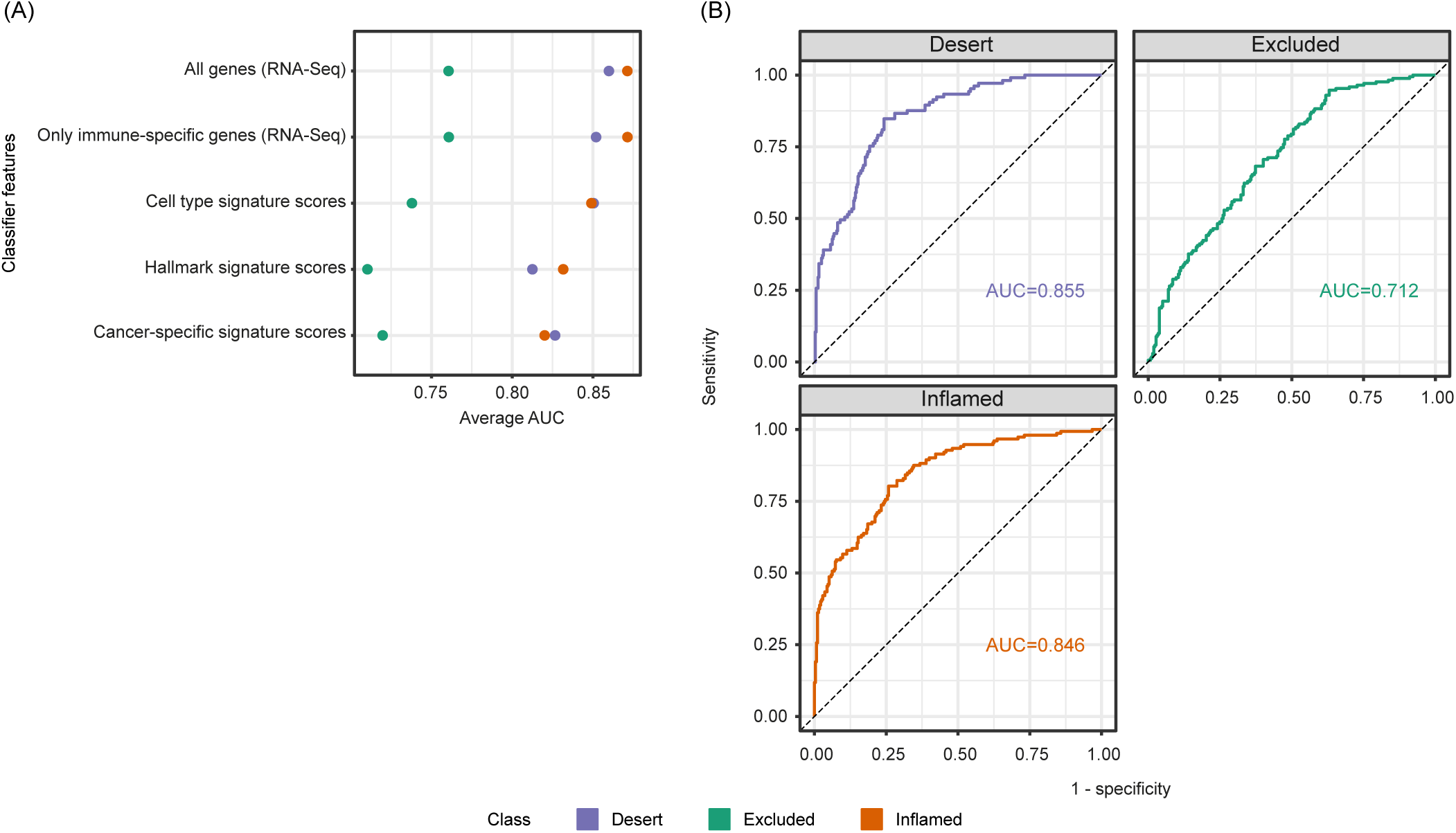
Accurate transcriptome-based prediction of CD8 immunophenotypes. (A) Predictiveness of genes and signatures: the performance of classifiers trained on distinct features, as assessed using 5-fold cross validation. (B) AUC curves for the final 92 gene-based classifier for CD8-desert, CD8-excluded, and CD8-inflamed phenotypes computed on the test dataset. AUC, area under receiver operating characteristic (ROC) the curve; RNA-seq, RNA-sequencing.

As the model trained on all genes had shown the best performance overall, we trained a final classifier based on the full transcriptome. A total of 92 genes were retained (supplemental figure S4B) in the classifier, which showed an AUC 0.855 for CD8-desert, 0.846 for CD8-inflamed, and 0.712 for CD8-excluded samples on the test dataset (**figure 4B, and methods**). Only 17% of the retained genes were highly correlated with CD8 expression and showed the pattern of high expression in inflamed, intermediate in excluded, and low expression in desert samples described in the exploratory section above (**supplemental figure S4C**).

### Both CD8+ T effector-associated and non-immune-expressed genes contribute to accurate immunophenotype classification

In order to explore, in detail, the characteristics of genes critically contributing to a good separation of the immunophenotype classes, we focused on genes retained in 85% of classifiers retrained on random subsets of the training data (**figures 5A–B and supplemental table S5**).

**Figure 5.**
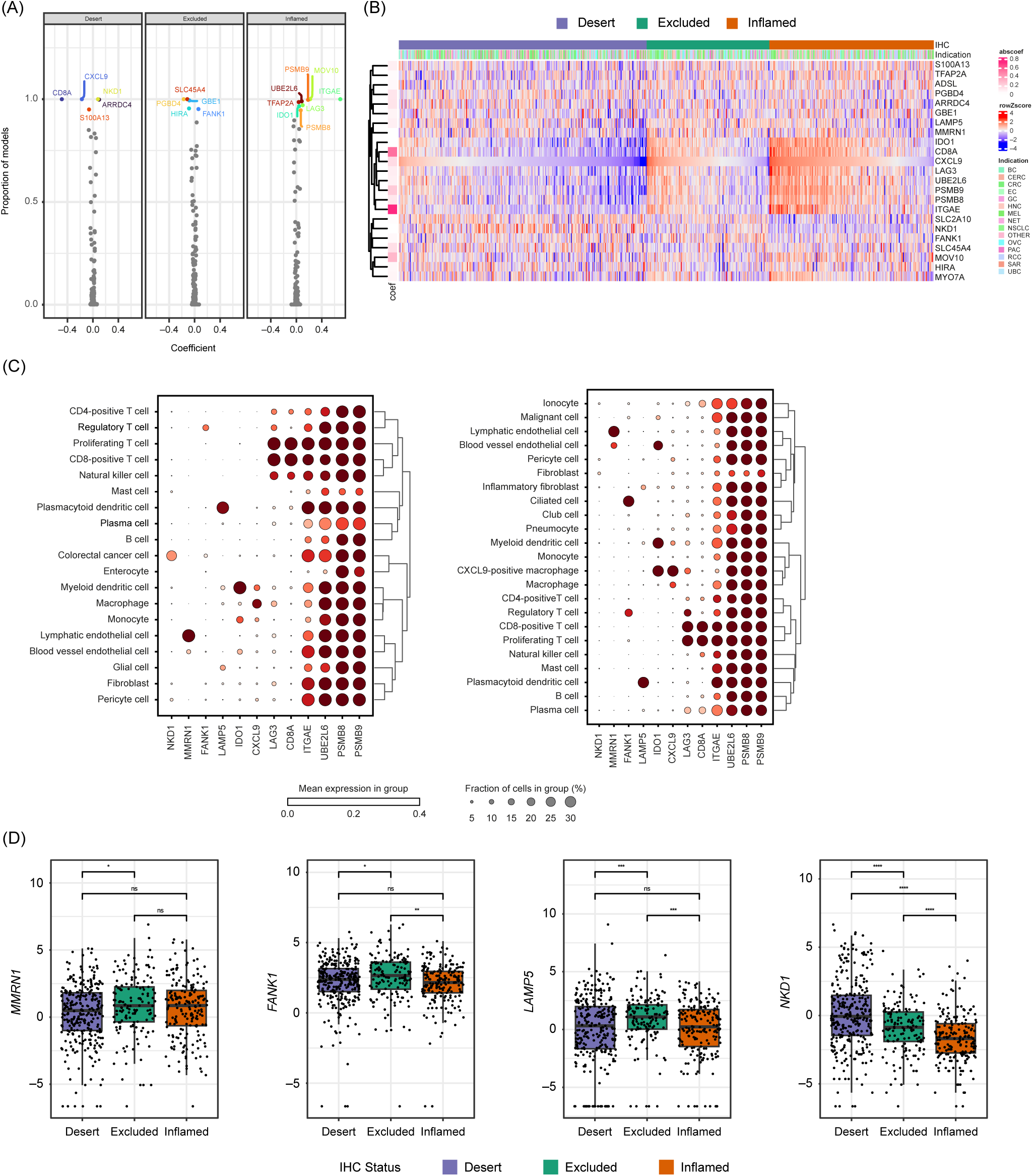
Characteristics of genes contributing highly to an accurate CD8 immunophenotype classification. (A) The frequency and magnitude of every feature in the RNA-seq classifier was measured by training and validating a classifier 100 times on random subsets of the data. The frequency of every feature and its median magnitude are displayed. Genes used in > 85% of the classifiers are labeled. (B) Heatmap showing the relative expression of top classification-relevant genes across CD8 immunophenotype class cancer types as well as in (C) CRC and lung cancer single-cell RNA-seq data and (D) bulk RNA-seq data. CRC, colorectal cancer; IgA, immunoglobulin A; IgG, immunoglobulin G; IHC, immunohistochemistry; LN, lymph nodes; mets, metastases; ns, not significant; RNA-seq, RNA-sequencing.

We examined their expression in bulk (this study) and in publicly available colorectal, lung and liver cancer single-cell RNA-seq data.^40–42^ The CD8+ T-effector/cytotoxicity, IFN-ɣ and antigen-processing/MHC pathway associated genes *IDO1, CD8A, CXCL9, LAG3, UBE2L6, PSMB8/9,* and *ITGAE* formed a cohesive cluster with increasing expression in desert-to excluded-to-inflamed samples (**figure 5B** **and supplemental figures S4C, S5A**). Despite this strong correlation, according to single-cell RNA-seq data, not all genes were expressed by the same cell subsets. *IDO1* and *CXCL9* were primarily expressed in the myeloid compartment, specifically on dendritic cells and macrophages (**figure 5C, supplemental figure S5B and supplemental table S5**). *LAG3, CD8A,* and *ITGAE* were CD8+ T-cell specific/enriched, while *UBE2L6, PSMB8,* and *PSMB9* showed ubiquitous expression, even beyond immune cell types (**figure 5C, supplemental figures S4C, S5B**).

The classifier also retained genes with opposite expression patterns (enriched in desert or enriched in excluded), some of which also showed highly cell-type-specific/enriched expression (**figure 5B** **and supplemental figure S5A–B**). For instance, the excluded-enriched *LAMP5* gene was mainly expressed in plasmacytoid dendritic cells. *FANK1* in regulatory T cells and malignant cells, while *MMRN1* showed endothelial-restricted expression (**figures 5B–D and supplemental figure S5B**). Among desert-enriched genes, we retrieved the glucose transported GLUT10 (encoded by *SLC2A10*) and the Wnt pathway inhibitor *NKD1,* which showed preferential expression in tumor cells and fibroblasts (**figures 5B–D and supplemental figure S5B**). By using publicly available spatial transcriptomics data in CRC (Valdeolivas et al. 2023^44^ and Wu et al. 2022^43^), we confirmed an enrichment in *NKD1* expression in CD8-desert as compared with CD8-high tumor areas. In addition, we observed the strongest expression of *LAMP5* in the sample showing CD8 infiltration in the stromal area only, in line with *LAMP5* being excluded-enriched (**supplemental figure S5C**).

In the training dataset, classifier features usually followed either desert-excluded-inflamed or inflamed-excluded-desert low-to-high expression patterns (**supplemental figure S4A, right columns of the plots**). To assess relationships between immunophenotypes on the molecular level, we performed uniform manifold approximation and projection (UMAP) transformation on the 92 genes used by the classifier trained on the gene expression values. We observed a clear separation between inflamed and desert, whereas excluded samples were more scattered (**supplemental figure S5D**). This suggests that the excluded immunophenotype is an intermediate state, rather than a distinct one. This is also consistent with lower performance of the classifiers when predicting the excluded phenotype.

### CD8 immunophenotype predictions are associated with patient survival and immunotherapy response

The developed classifier was applied to the pan-cancer TCGA dataset in order to evaluate its prognostic effect on OS. The first step towards this was to batch-correct the TCGA dataset using only lung samples as a reference. The phenotypes of hematologic malignancies, diffuse large B-cell lymphoma, and thymoma were predicted as CD8-inflamed only (**figure 6A**). Classically highly inflamed indications including melanoma, lung squamous cell carcinoma, and RCC were predicted to have ≥ 50% CD8-inflamed samples, compared with tumors with typically low levels of inflammation, such as pancreatic and prostate cancer, where only < 10% of samples were predicted as CD8-inflamed.

**Figure 6.**
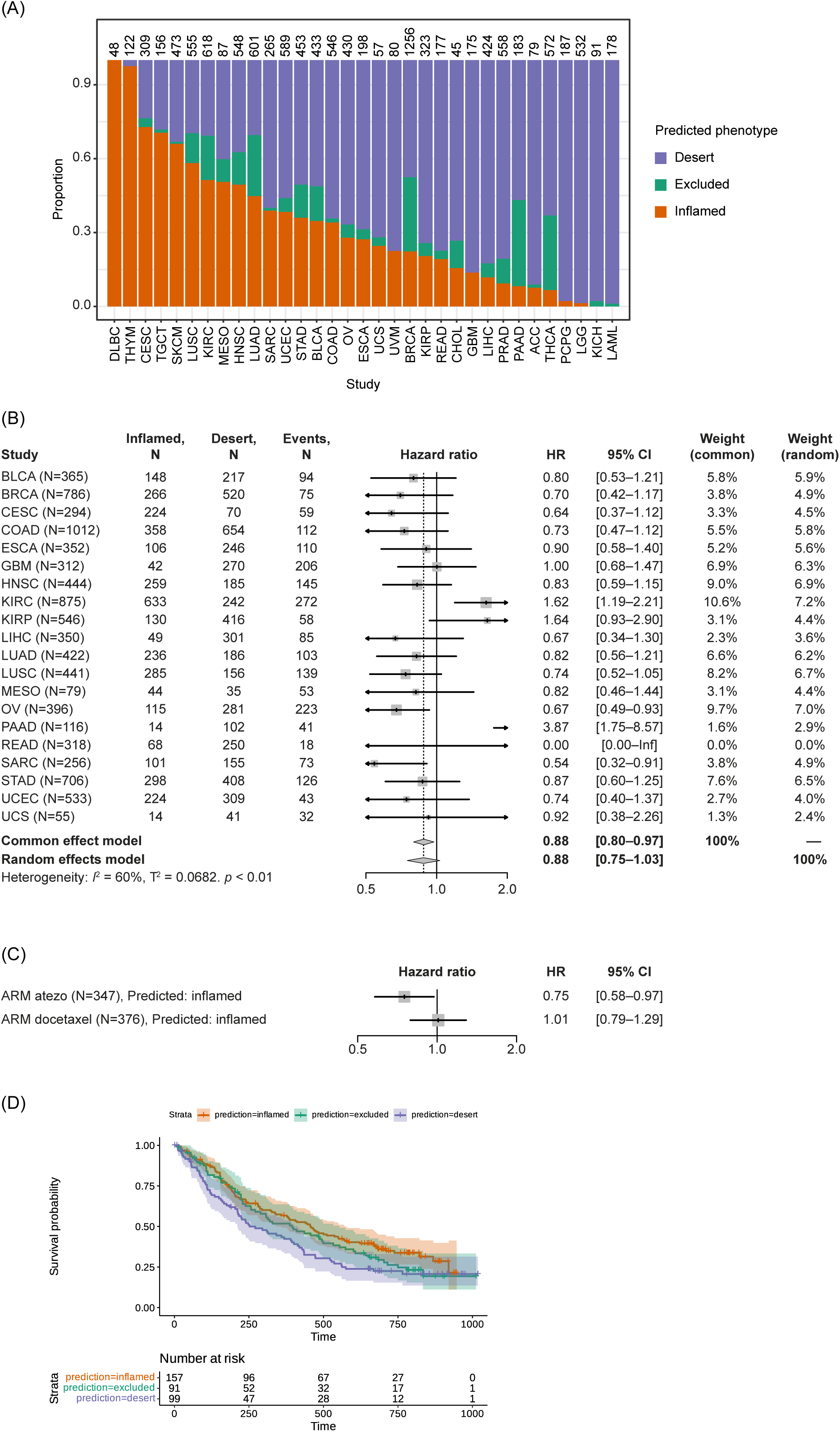
Application of the transcriptome-based classifier on patient data. (A) Predicted CD8 immunophenotypes across TCGA. (B) Prognostic value of predicted CD8 inflamed versus desert immunophenotype based on TCGA data. Samples predicted as excluded were removed from the analysis. (C) Predictive value of the classifier in lung cancer. CI, confidence interval; CIT, cancer immunotherapy; HR, hazard ratio; TCGA, The Cancer Genome Atlas.

A positive association was observed between the predicted CD8 immunophenotype and OS across indications. Specifically, patients who had predicted CD8-inflamed tumors across TCGA showed prolonged OS compared with patients who had CD8-desert tumors (hazard ratio [HR] 0.88; 95% confidence interval [CI]: 0.80–0.97) (**figure 6B**). While ovarian carcinoma and sarcoma showed particularly strong survival benefits, kidney cancer was the sole indication associated with a worse prognosis in patients with inflamed phenotypes (**figure 6B**).

Finally, the classifier was evaluated using data from the OAK trial, including patients for whom the IHC-based CD8 immunophenotype was missing. In the atezolizumab arm, samples predicted as inflamed had a HR for mortality of 0.75 (95% CI: 0.58–0.97), whereas in the docetaxel arm, the HR was 1.01 (95% CI: 0.79–1.29), suggesting the CD8 immunophenotypes only had an influence on the survival of immune checkpoint inhibitor-treated patients and not on patients treated with docetaxel (**figure 6C**).

Finally, we investigated if newly identified immunophenotype-enriched genes showed any association with increased/decreased survival across TCGA. While no significant patterns were observed for *NKD1*, *LAMP5*-high tumors showed a higher risk of mortality compared with the overall tumor samples: the common effect model HR for mortality was 1.20 (95% CI: 1.08– 1.33) (**supplemental figures S6A–C**).

## DISCUSSION

IHC analyses of CD8+ T-cell infiltration in tumor samples represents one widely used approach to stratify patients by immunophenotype based on the spatial location of CD8+ T cells in the TME. To better understand variability in this phenotype across indications and excision locations, as well as to what extent is it reflected in transcriptional patterns, we generated and explored a large dataset (> 2000 samples) from 14 phase I–III trials, in which the IHC data were scored and classified in an identical way.

Consistent with prior reports, we found that CD8 immunophenotypes showed highly distinct patterns between indications as well as between primary and metastatic sites.^12^ Strikingly, CD8-deserts were most abundant in liver metastases, regardless of the tumor origin. Together with the fact that the majority of biopsies in our phase I/II cohort originated from liver metastases, despite studies not normally mandating exact biopsy locations, this highlights that the patient population enrolled in CIT trials may be highly challenging to treat due to low baseline immune infiltration. In contrast, indications such as melanoma, lung and bladder cancers presented a predominant CD8-inflamed and/or CD8-excluded phenotype, in line with these tumors generally being considered to be immune ‘hot’ and responsive to CIT.^47 48^ Our findings highlight the impact of the local microenvironment on tumor immunophenotype as well as on biomarker or pharmacodynamics analyses and treatment decisions based on biopsies from metastases.

Inflammatory and suppressive immune signatures also varied in areas of differential CD8 immunophenotypes. The strongest signal was related to T-effector and IFN-ɣ activity, which showed highest levels in inflamed, intermediate in excluded and very low levels in desert samples, in line with high consistency at protein and RNA level. Few excluded-specific signals were detected: 45 genes showed significantly higher expression in both excluded versus deserts and excluded versus inflamed comparisons, enriching for fibroblast, endothelial and mast cell-specific expression. Interestingly, mast cells can take multiple roles in shaping the TME, promote angiogenesis and support tumor invasiveness,^49^ and have recently been shown to mediate resistance to anti-programmed death-1 (anti-PD1) therapy.^50^ Further, the excluded-enriched expression of stromal-specific genes and the angiogenic and TGF-β pathways is consistent with an immunosuppressive environment, underlying immune cell exclusion.^2^ Our findings are also in line with a recent ovarian cancer study, which found that CD8-excluded phenotypes have a higher stromal, TGF-β, and angiogenesis pathway activity.^22^

The good correspondence between the transcriptome and IHC-based measurements encouraged us to develop a classifier that can accurately predict IHC-derived CD8 immunophenotypes solely based on gene expression.

Our classifier yielded a high predictive performance, albeit better for CD8-inflamed/CD8-desert phenotypes than for the CD8-excluded phenotype. This lower performance for the excluded phenotype may be because RNA-seq provides a bulk characterization of the tumor, while the excluded phenotype is largely characterized by spatial relationships more difficult to capture at the molecular level. Notably, the described CD8 IHC immunophenotype classification relies on a pathologist’s assessment of lymphocyte location on a whole-slide tissue with regards to tumor epithelium and stroma areas, which makes the method prone to interobserver bias. Given this inherent noise, the performance of the classifier was very high.

We tested the broad applicability of the classifier by evaluating the prognostic effect on overall survival (OS) in pan-cancer TCGA and clinical trial datasets, and the predictive effect on OS in lung cancer from the OAK study. Both the content of CD8+ T cells and many of the pathways and signatures identified and described in this report were previously found to be positively associated with survival and response to cancer immunotherapy.^3 10 13 51 52^ For instance, *CXCL9* and *CD8A* (two of our top classifier genes), as well as the inflamed signature,^53^ which includes *CD8A*, *LAG3* and *IDO1* from our top classifier genes, were positively associated with CIT response according to a recent meta-analysis across > 1000 patients.^10^ Consistent with these observations, patients with NSCLC in the OAK trial^26^ receiving atezolizumab had superior OS with classifier-predicted CD8-inflamed tumors than with CD8-desert tumors, whereas there was no OS difference between those phenotypes in those patients receiving docetaxel.

When investigating the prognostic effect of the predicted CD8 immunophenotype across the TCGA data, a positive association was found between CD8 immunophenotype and OS across indications, as expected based on previous reports.^54^ Patients with kidney cancer with inflamed phenotypes had a worse prognosis than those with other CD8 immunophenotypes. This atypical pattern of RCC was recently reported in a study where the proliferating CD8+ T cell percentage was histologically assessed.^55^ Thus, our predictions are in line with previous findings but expand the range of indications for which the relation between immunophenotype and response can be evaluated.

While most of the T-effector and immune infiltration-related signals have been previously identified and described, our approach also revealed predictive features stemming from other cellular compartments. This included the desert-enriched CRC gene *NKD1*, which is induced to antagonize Wnt signaling and promotes cancer cell proliferation.^56^ Although we confirmed the tumor-associated localization of *NKD1* using publicly available spatial transcriptomic data, we did not detect a significant survival association across TCGA data. However, we report here for the first time that excluded-enriched *LAMP5* expression is negatively associated with OS across indications in TCGA data. Interestingly, *LAMP5* depletion was recently found to significantly inhibit leukemia cell growth and to be a modulator of innate immune pathways by suppressing type I IFNs downstream of TLR9.^57 58^ *LAMP5* was also detected on TGF-β-myofibroblastic cancer-associated fibroblasts, which are associated with an immunosuppressive environment.^59^ Our single-cell RNA-seq analysis confirms fibroblasts, along with plasmacytoid dendritic cells, as the main source of *LAMP5* expression in tumor tissues. How LAMP5 contributes to immune cell exclusion and a survival disadvantage, and which of the two cell types is important for this association, warrants further investigation.

Potential limitations of our study include the need to further confirm our findings in a wider population and other cancer types. Notably, our RNA-seq-based classifier was solely developed to predict the CD8 immunophenotype, and has thus not been optimized to predict survival benefit, in contrast to other studies highlighting CD8-related biomarkers. We solely reported here a correlation between the predicted CD8 immunophenotype and OS. Finally, the bulk RNA-seq method used in our study has previously shown limited reproducibility due to heterogeneity between tumor cells and within the microenvironment.^60^ More recent technologies such as single-cell RNA-seq and spatial transcriptomics could address the limitations of bulk RNA-seq and provide more accurate information such as spatial resolution.^2 60^

## CONCLUSIONS

CD8 immunophenotyping has emerged as a powerful tool for understanding the TME and its impact on cancer. Our novel 92-gene classifier accurately predicts the spatial CD8 immunophenotype of primary and metastatic tumors. As RNA-seq provides wider information on patient samples than IHC, the new classifier could be used for retrospective and reverse translation analyses of CD8 immunophenotypes from clinical trial cohorts without the need for individual tissue section curation.

Tumor-agnostic enrichment strategies require consideration of spatial location of immune cells, immune-related patterns and lesion location,^61^ and the development of this RNA-based CD8 immunophenotyping classifier is a promising step in this direction, providing a reliable, cost-effective and simple tool to help optimize patient selection, response to cancer immunotherapy, and patient outcomes.

## Declarations

### Ethics approval and consent to participate

All clinical studies were conducted in accordance with the principles of the Declaration of Helsinki and Good Clinical Practice Guidelines. Written informed consent was collected from all enrolled patients.

### Consent for publication

Not applicable.

### Availability of data and material

The source code used for the analysis is available via: https://github.com/bedapub/CD8-immune-phenotype-paper-supplementary. The classifier is available as an R-package at: https://github.com/bedapub/cd8ippred. For up to date details on Roche’s Global Policy on the Sharing of Clinical Information and how to request access to related clinical study documents, see here: https://go.roche.com/data_sharing. The data may be shared in a pseudonymized way. Anonymized records for individual patients across more than one data source external to Roche can not, and should not, be linked due to a potential increase in risk of patient re-identification.

### Competing interests

AR, IID, PCS, MS-S, NS, AV, KK are employees of F. Hoffmann-La Roche Ltd. AH, CSF, GD, MC are employees of Roche Diagnostics GmbH. IK was an employee of Roche Diagnostics GmbH at the time of the study. AH, AR, IID, PCS, MS-S, NS, CSF, AV, KK, MC are shareholders of F. Hoffmann-La Roche Ltd.

### Funding

This study was supported by F. Hoffmann-La Roche Ltd.

### Author contributions

AR, IID, PCS, MS-S and MC designed the study, interpreted the data and drafted the manuscript, with additional input from all authors. IID designed and executed the machine learning experiment, including classifier training, validation, assessment of feature importance, developed and executed batch-correction methodology, and performed survival analyses. PCS performed analyses relating to signature enrichment, gene expression characteristics in bulk, and single-cell transcriptomics data. AV performed analyses related to spatial transcriptomics data. MS-S performed analyses relating to cohort characterization. AH performed cross-clinical trial sample metadata curation. AH and CSF were responsible for CD8IHC immunophenotype data integration and harmonization. NS performed the differential expression analyses. All authors read and approved the final manuscript.

## Acknowledgments

The authors would like to thank the patients, their families, and the participating study centers. We acknowledge Laura Jarassier for data analysis support and the Roche-wide enhanced Data and Insights Sharing (EDIS) network for providing curated and annotated clinical data. Medical writing support for the development of this manuscript, under the direction of the authors, was provided by Laura Vergoz, PhD, and Edward Neale, PhD, of Ashfield MedComms, an Inizio company, and funded by F. Hoffmann-La Roche Ltd.

## Abbreviations

AUC: area under the curve
CI: confidence interval
CIT: cancer immunotherapy
CRC: colorectal cancer
FDR: false discovery rate
HR: hazard ratio
IE: intra-epithelial
IFN: interferon
IHC: immunohistochemistry
ITS: intra-tumoral stroma
MSigDB: Molecular Signature Database
NSCLC: non-small-cell lung cancer
OS: overall survival
PCA: principal component analysis
PD-1: programmed death 1
RCC: renal cell carcinoma
RNA-seq: RNA-sequencing
RT-PCR: real-time polymerase chain reaction
TCGA: The Cancer Genome Atlas
TME: tumor microenvironment
UMAP: uniform manifold approximation and projection.

## Notes

https://github.com/bedapub/cd8ippred

https://github.com/bedapub/CD8-immune-phenotype-paper-supplementary

