## Supplementary Figures for "Tumor-agnostic transcriptome-based classifier identifies spatial infiltration patterns of CD8+ T cells in the tumor microenvironment and predicts clinical outcome in early- and late-phase clinical trials"

Figure S1

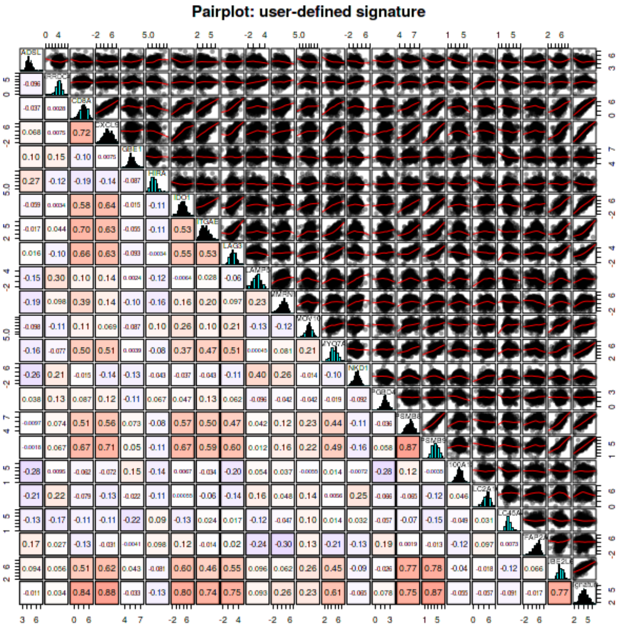


**Figure S1** Pairwise correlation between all (92) classification-relevant genes across the training dataset.

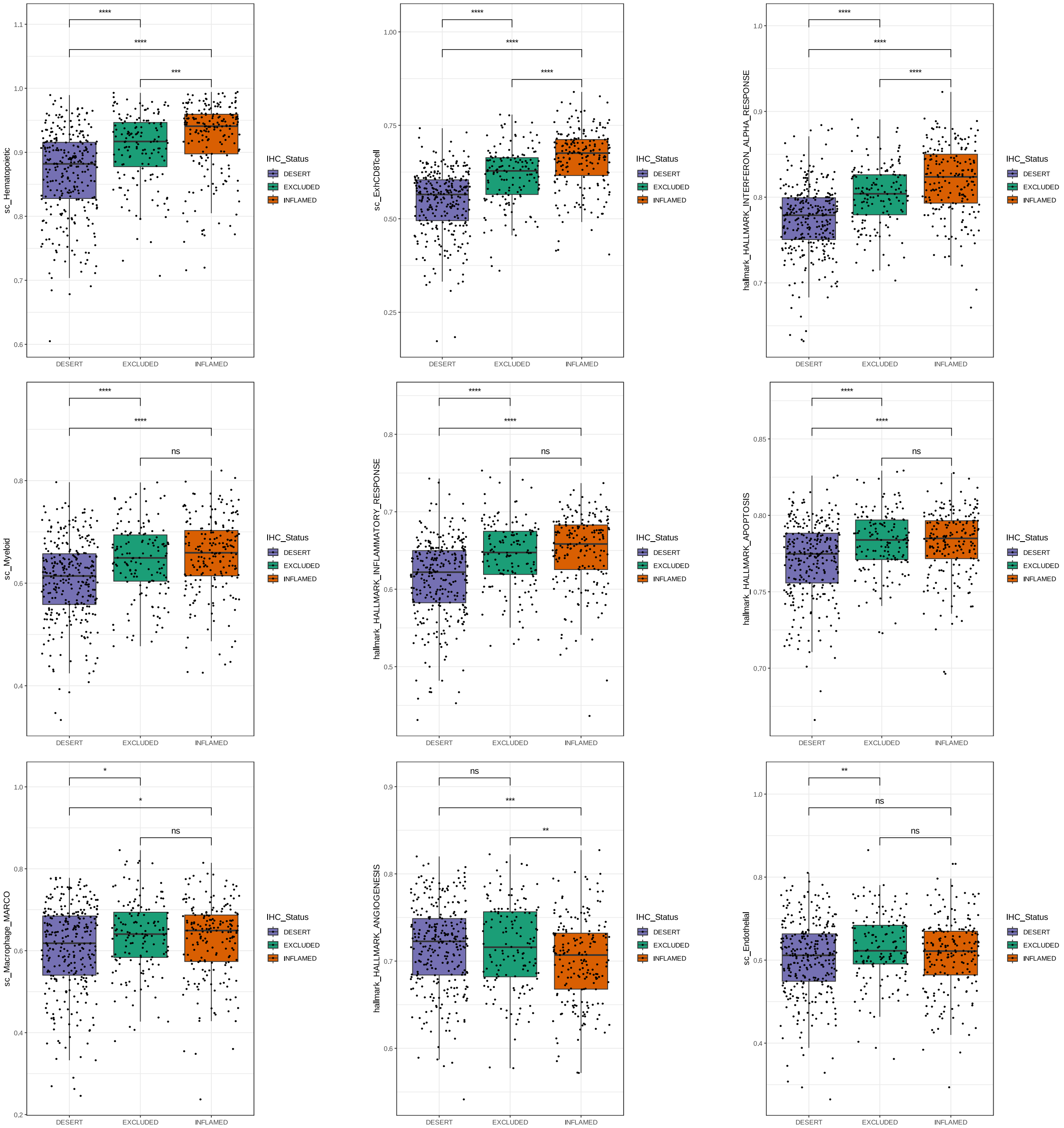


Figure S2 Gene expression related to CD8 immunophenotypes. IHC, immunohistochemistry.

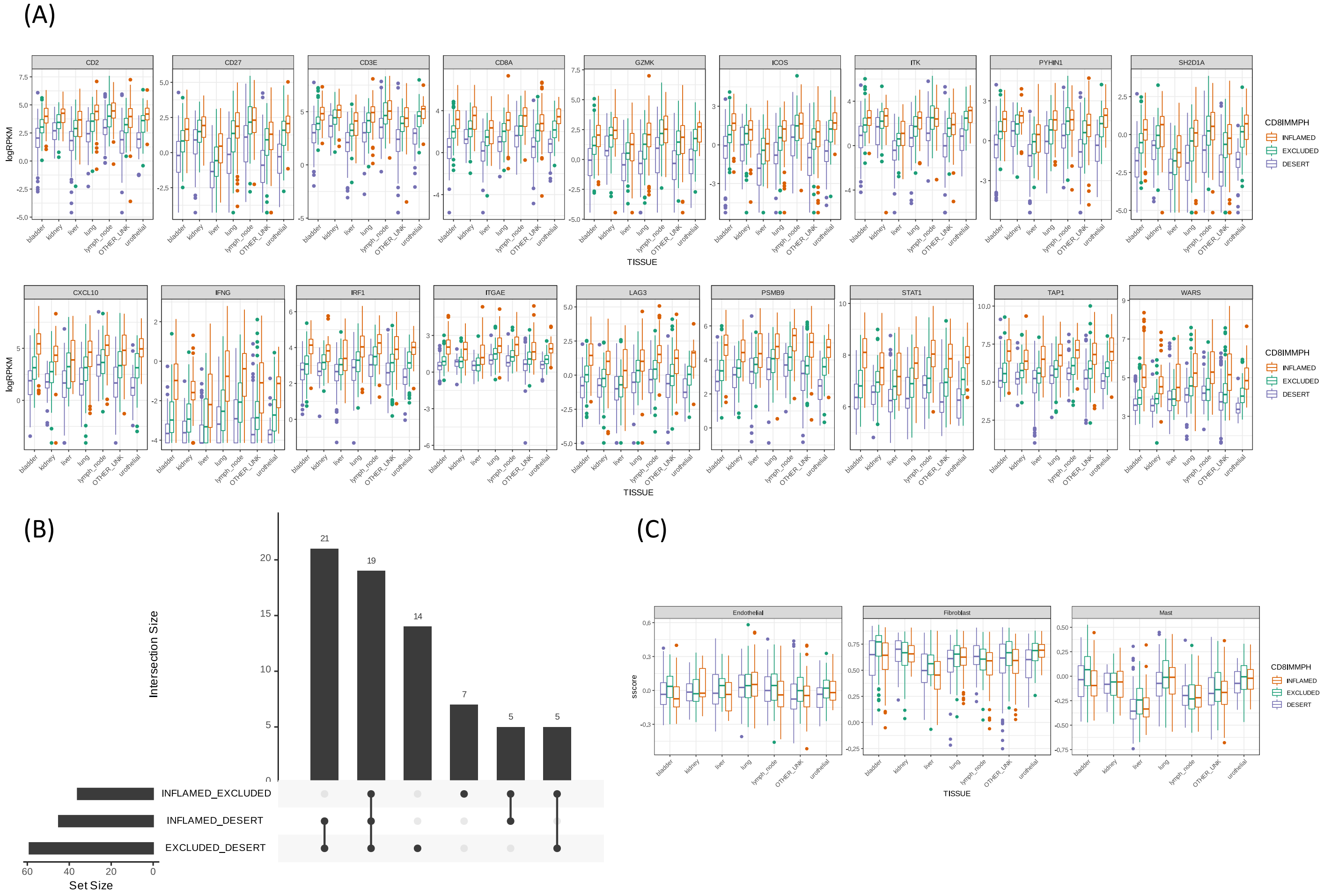


**Figure S3** Systematic characterization of gene expression differences across CD8 immunophenotype classes. (A) Overview of expression distribution for top differentially expressed genes across pairwise comparisons, stratified by excision site. (B) Overlaps between the signatures significantly differentially enriched across all pairwise comparisons. (C) Enrichment (rank biserial correlation) of CD8-excluded associated signatures across CD8 immunophenotype classes and excision sites. CD8IMMPH, CD8 immunophenotype; logRPKM, log Reads Per Kilobase of transcript, per Million mapped reads.

Figure S4

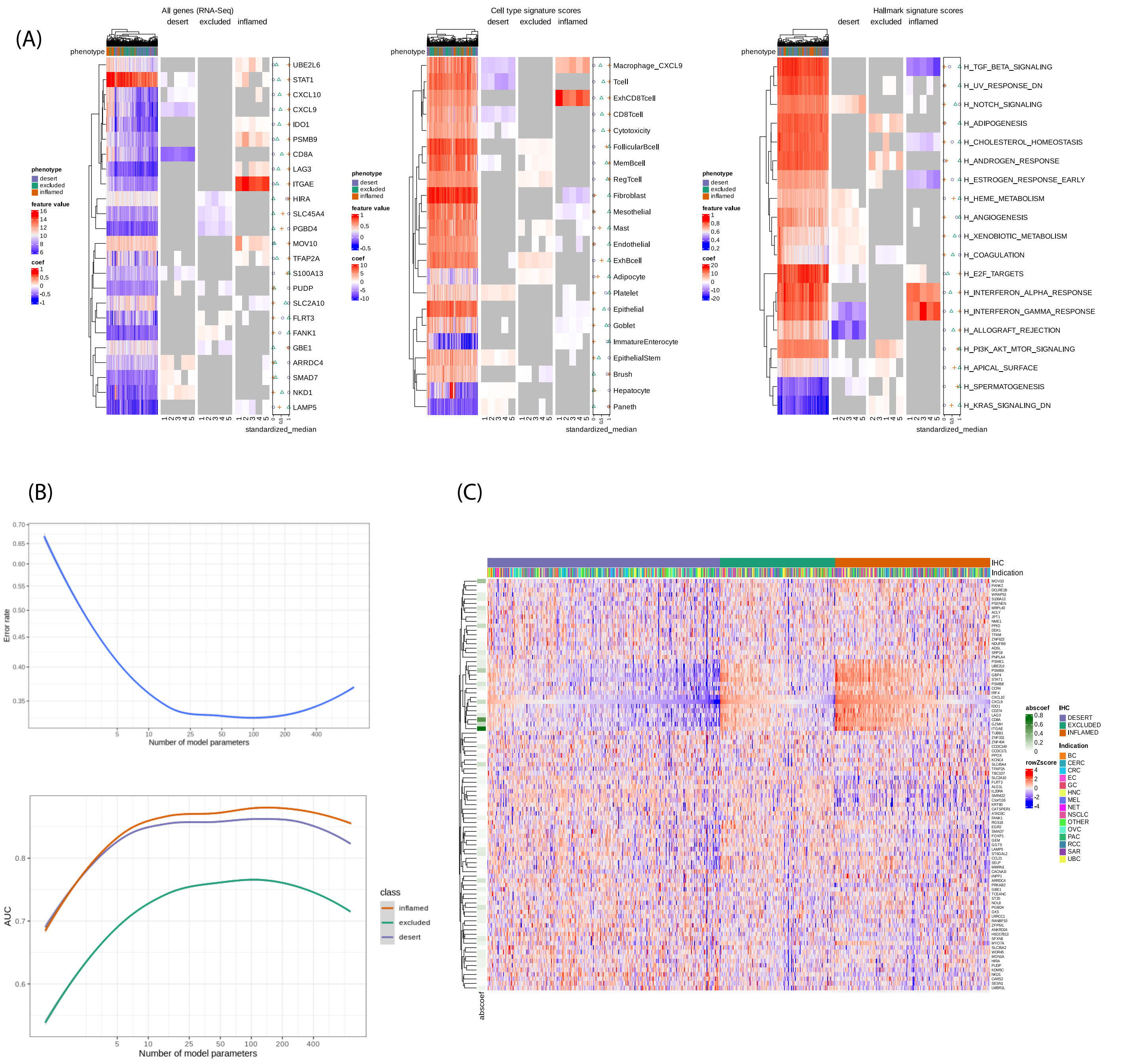


Figure S4 Development of a transcriptomebased CD8 immunophenotype classifier. (A) Genes and signatures used by distinct transcriptomebased classifiers, trained on (left) all genes, (middle) cell type specific signature scores and (right) hallmark pathway scores. A 5fold cross-validation across the training data was used and only features present at least in four iterations are shown. The left section of every tile shows expression among the training samples (columns). The middle section shows the classifier coefficient in every iteration. Gray color indicates that a feature was not used in that iteration. The right section shows relative median expression value among the CD8 immunophenotypes. (B) Performance of the classifier depending on the number of features used, assessed using inner cross validation. (C) Heatmap showing the relative expression of all (92) classificationrelevant genes across CD8 immunophenotype classes, tumor indications and excision sites.

### Figure S5A & B

(A)

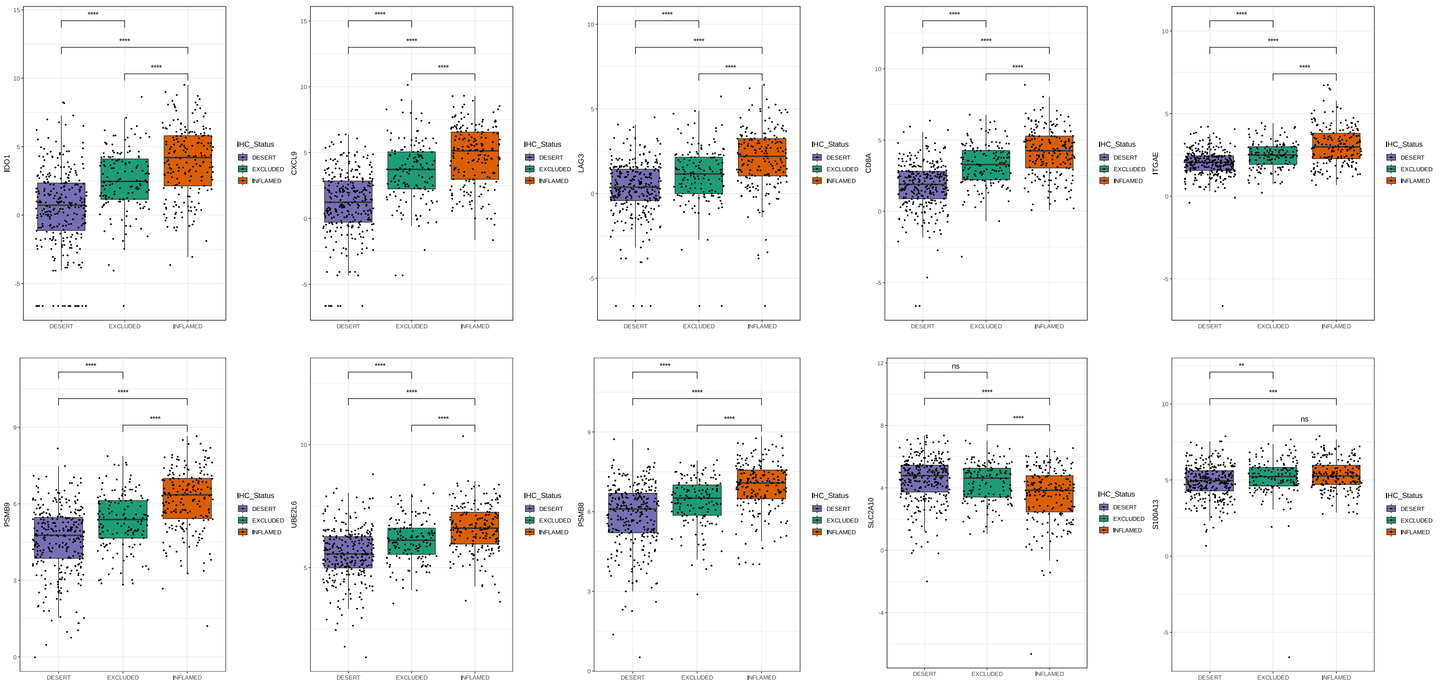


(B)

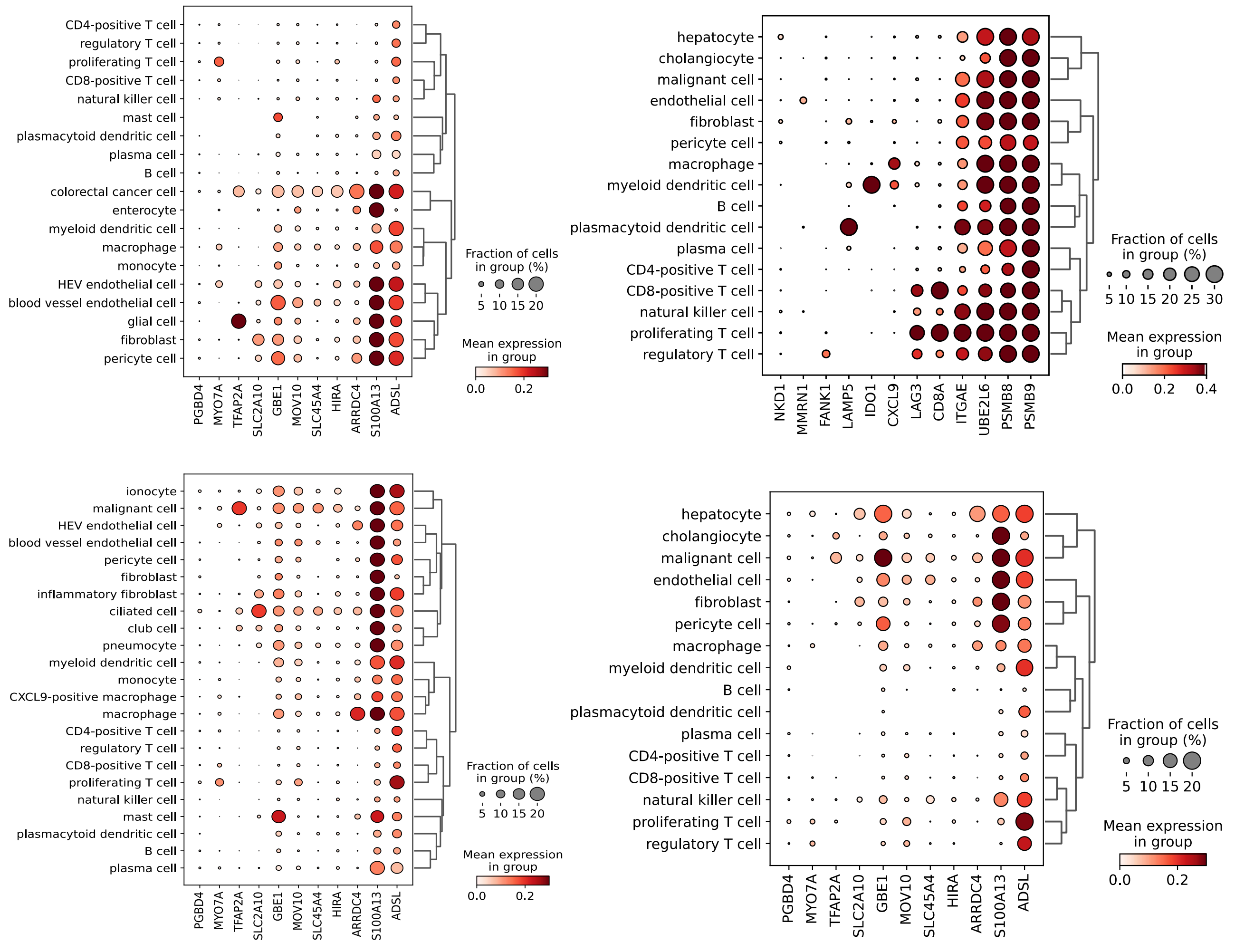

(C)

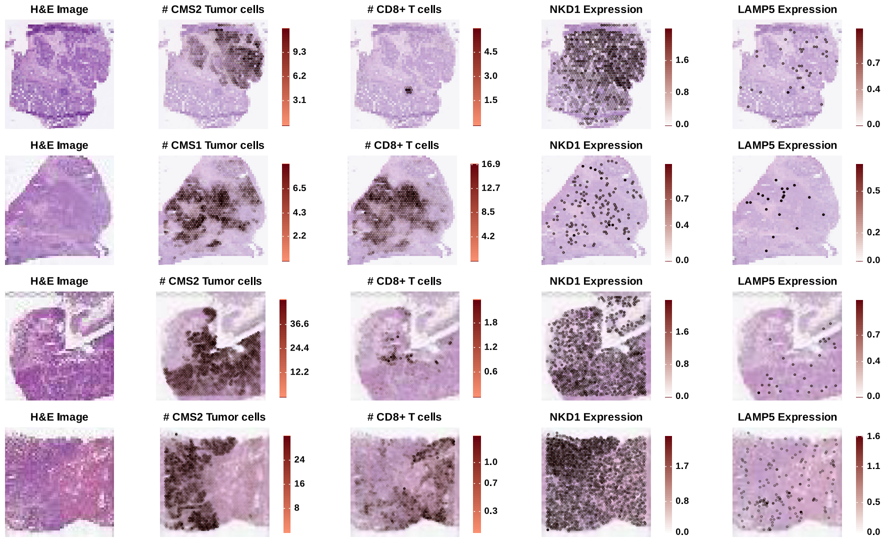


(D)

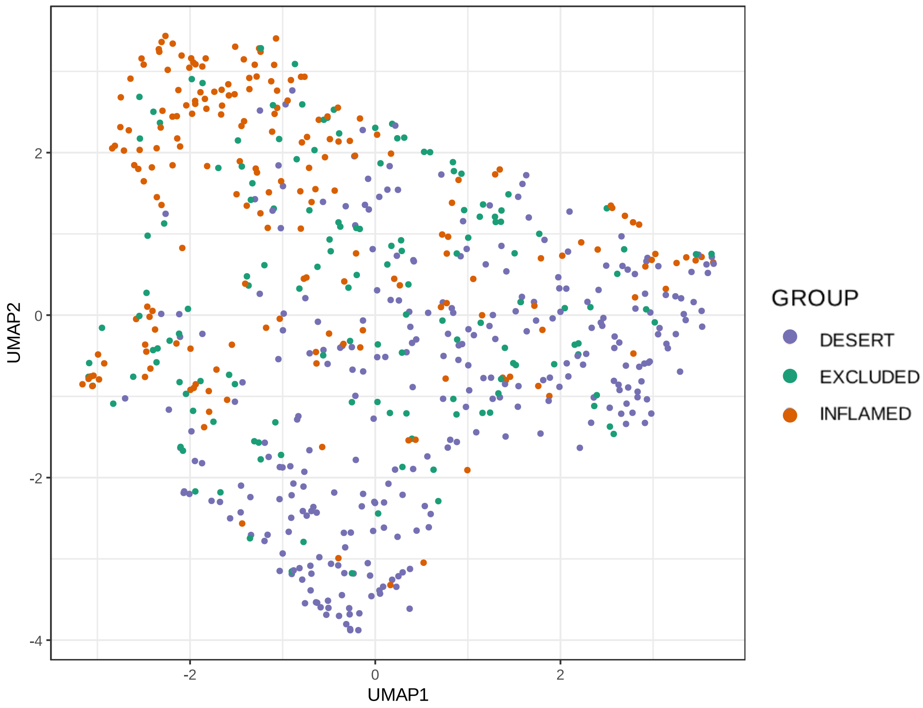


Figure S5 Characteristics of genes highly contributing to an accurate CD8 immunophenotype classification. (A) Expression of selected CD8-inflammed associated genes in bulk RNA-seq data. (B) Expression of CD8-immunophenotype-enriched genes in colorectal cancer (CRC), lung cancer and liver cancer single-cell RNA-seq data, stratified per cell type. (C) Gene expression maps of NKD1 and LAMP5 in selected spatial transcriptomics samples derived from CRC patients. CMS1 tumors are known to present large infiltration of CD8+ T cells, while CMS2 tumors are considered an immune-desert. The bottom sample corresponds to liver metastases of a primary CRC tumor that is shown immediately above it. (D) Samples from the training dataset in a 2D UMAP space generated based on the expression values of all (92) genes selected by the classifier. IHC, immunohistochemistry.

(A)

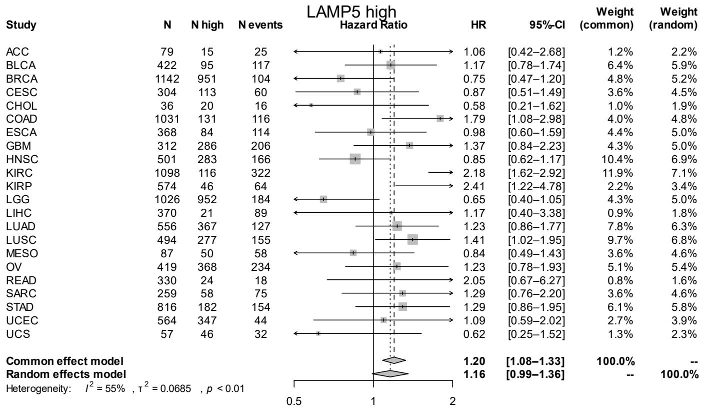


(B)

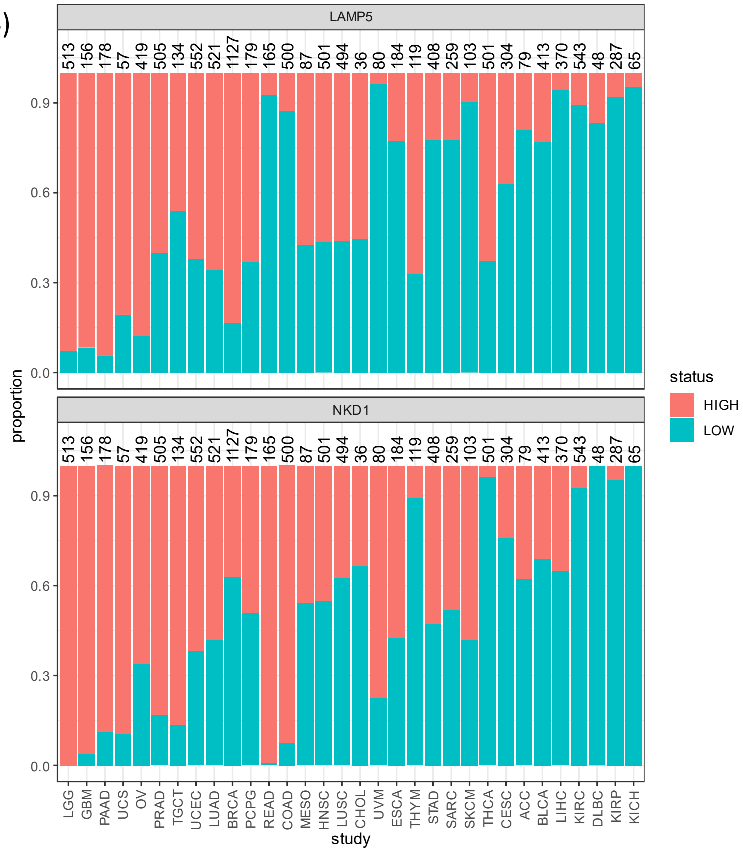


(C)

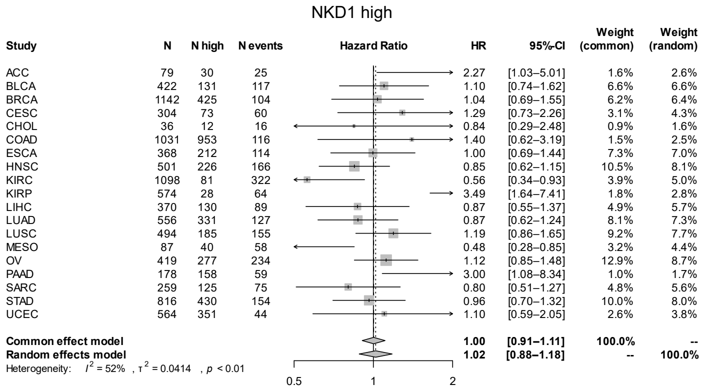


**Figure S6** Survival association of novel genes contributing to CD8 immunophenotype classification. (A) Hazard ratio for mortality in *LAMP5*-high tumors vs all tumors. (B) Status of *LAMP5* and *NKD1* expression in various TCGA indications. (C) Hazard ratio for mortality in *NKD1*-high tumors vs all tumors. All study names and codes can be found on TCGA The Cancer Genome Atlas's portal at https://gdc.cancer.gov/resources-tcga-users/tcga-code-tables.
