## Supplementary Tables for "Tumor-agnostic transcriptome-based classifier identifies spatial infiltration patterns of CD8+ T cells in the tumor microenvironment and predicts clinical outcome in early- and late-phase clinical trials"

**Table S1** List of phase I/II clinical trials from which tissue samples were sourced.

| **NCT number** | **Phase** | **Full trial name** | **Tumor type(s)** | **Drug(s)** | **No of samples used in this analysis** |
| --- | --- | --- | --- | --- | --- |
| **NCT02004106** | I | A Study to Evaluate Safety, Pharmacokinetics, and Efficacy of RO6895882 in Participants With Advanced and/or Metastatic Solid Tumors | Neoplasms | RO6895882 | 64 |
| **NCT02304393** | I | A Study of Selicrelumab (RO7009789) in Combination With Atezolizumab in Participants With Locally Advanced and/or Metastatic Solid Tumors | Solid tumors | Atezolizumab; Selicrelumab | 91 |
| **NCT02323191** | I | A Study of Emactuzumab and Atezolizumab Administered in Combination in Participants With Advanced Solid Tumors | Solid cancers | Atezolizumab; Emactuzumab | 138 |
| **NCT02350673** | I | A Study of Intravenous (IV) Cergutuzumab Amunaleukin and Atezolizumab in Combination in Participants With Locally Advanced and/or Metastatic Solid Tumors | Solid tumors | Atezolizumab; Cergutuzumab Amunaleukin | 36 |
| **NCT02627274** | I | A Study Evaluating Safety, Pharmacokinetics, and Therapeutic Activity of RO6874281 as a Single Agent (Part A) or in Combination With Trastuzumab or Cetuximab (Part B or C) | Solid tumors;  Breast cancer;  Head & Neck cancer | RO6874281; Trastuzumab; Cetuximab | 57 |
| **NCT02665416** | I | Study Evaluating the Safety, Pharmacokinetics (PK), Pharmacodynamics (PD), and Therapeutic Activity of Selicrelumab (RO7009789) With Vanucizumab or Bevacizumab in Participants With Metastatic Solid Tumors | Advanced/metastatic solid tumors | Selicrelumab; Vanucizumab; Bevacizumab | 52 |
| **NCT03063762** | I | Study to Evaluate Safety, Pharmacokinetics and Therapeutic Activity of RO6874281 as a Combination Therapy in Participants With Unresectable Advanced and/or Metastatic Renal Cell Carcinoma (RCC) | Renal cell carcinoma | Atezolizumab; Bevacizumab; RO6874281 | 46 |
| **NCT03539484** | I | A Study of RO7172508 in Patients With Locally Advanced and/or Metastatic CEA-Positive Solid Tumors | Solid tumors | RO7172508; Obinutuzumab; Tocilizumab | 16 |
| **NCT03292172** | I | A Study to Evaluate the Safety, Pharmacokinetics and Clinical Activity of RO6870810 and Atezolizumab (PD-L1 Antibody) in Participants With Advanced Ovarian Cancer or Triple Negative Breast Cancer | Advanced ovarian cancer; Triple negative breast cancer | Atezolizumab; RO6870810 | 17 |
| **NCT03386721** | II | Basket Study to Evaluate the Therapeutic Activity of Simlukafusp Alfa as a Combination Therapy in Participants With Advanced and/or Metastatic Solid Tumors | Advanced/ metastatic Head & Neck, cervical and esophageal cancers | Simlukafusp; Atezolizumab; Gemcitabine; Vinorelbine | 90 |
| **NCT02031458** | II | A Study of Atezolizumab in Participants With Programmed Death - Ligand 1 (PD-L1) Positive Locally Advanced or Metastatic Non-Small Cell Lung Cancer (BIRCH) | NSCLC | Atezolizumab | 21 |

Further information on these trials can be found at clinicaltrials.gov. NSCLC, non-small-cell lung cancer

**Table S2** List of differentially expressed genes. See separate Excel spreadsheet.

**Table S3** List of differentially expressed signatures. [See separate Excel spreadsheet](https://docs.google.com/spreadsheets/d/1fSluUNp7dn8eZq_P2CgZ1VsSatYLYNS2/edit#gid=24502773).

**Table S4** Patient baseline characteristics.

| Characteristic |  | Patient population (N=628) |
| --- | --- | --- |
| Age | Median (range) | 60 (18–86) |
| Sex, n (%) | Female | 281 |
|  | Male | 347 |
| Tumor sample origin*, n (%) | NSCLC (Non-small cell lung cancer) | 89 |
|  | CRC (Colorectal carcinoma) | 84 |
|  | HNC (Head and neck carcinoma) | 60 |
|  | RCC (Renal cell carcinoma) | 53 |
|  | UBC (Urinary bladder carcinoma) | 51 |
|  | OVC (Ovarian carcinoma) | 37 |
|  | BC (Breast carcinoma) | 33 |
|  | PAC (Pancreatic adenocarcinoma) | 32 |
|  | MEL (Melanoma) | 28 |
|  | EC (Esophageal carcinoma) | 28 |
|  | SAR (Sarcoma) | 17 |
|  | GC (Gastric carcinoma) | 14 |
|  | CERC (Cervical carcinoma) | 14 |
|  | NET (Neuroendocrine tumor) | 9 |
|  | AdrenC (Adrenocortical carcinoma) | 5 |
|  | Meso (Mesothelioma) | 4 |
|  | CholC (Cholangiocarcinoma) | 3 |
|  | IMEL (Intraocular Melanoma) | 3 |
|  | PRCA (Prostate cancer) | 3 |
|  | SIC (Small intestine carcinoma) | 3 |
|  | AnC (Anal carcinoma) | 2 |
|  | SkinC (Skin carcinoma) | 2 |
|  | ENDC (Endometrial carcinoma) | 1 |
|  | Thymo (Thymoma) | 1 |
|  | ThyrC (Thyroid carcinoma) | 1 |
| CD8 immune phenotype†, n (%) | Inflamed | 193 (30.7) |
|  | Desert | 144 (22.9) |
|  | Excluded | 291 (46.3) |
| Any previous therapy, n (%) | Experienced | 126 (20.1) |
|  | Naïve | 493 (78.5) |
|  | Missing | 9 (1.4) |
| Previous ICIs‡, n (%) | Lapcin + nivolumab | 59 (9.3) |
|  | Pembrolizumab | 29 (4.6) |
|  | Atezolizumab | 16 (25.5) |
|  | Other minor ones (avelumab, durvalumab, cemiplimab, ipilimumab) | 14 (2.2) |
|  | Combinations (ipilimumab- nivolumab, durvalumab, avelumab, ipilimumab-atezolizumab) | 8 (1.3) |

*Tumor types with n < 5 samples (25 different categories: eg, bone, fallopian tube, pelvis, thymus, testis; most of them n=1) are not included. The percentages have therefore not been calculated; †CD8-inflamed was defined as CD8/Ki67 counts > 500 cells/mm^2^ in the intraepithelial compartment with IE2 + IE3 ≥ 20%. CD8-excluded was defined as ≤ 500 cells/mm^2^ in the intraepithelial and > 50 cells/mm^2^ in the stromal compartment ITS0 + ITS1 ≥ 80% and ITS2 + ITS3 < 20%. CD8-desert was defined as ≤ 50 cells/mm^2^ in the stromal compartment IE0 + IE1 ≥ 80% & IE2 + IE3 < 20% and ITS0 + ITS1 < 80% and ITS2 + ITS3 ≥ 20%; ‡n=126 patients had prior therapy. ICI, immune checkpoint inhibitor.

**Table S5** Overview of the top genes identified from the machine-learning approach.

| **Name** | **Desert** | **Excluded** | **Inflamed** | **Tissue_sc_summary** |
| --- | --- | --- | --- | --- |
| IDO1 |  |  | up | cDC_(endothelial) |
| ITGAE |  |  | up | cytotox_enriched |
| LAG3 |  |  | up | CD8T |
| PSMB8 |  |  | up | broad_immunecor |
| PSMB9 |  |  | up | broad_immunecor |
| UBE2L6 |  |  | up | broad_immunecor |
| CD8A | down |  |  | CD8T |
| CXCL9 | down |  |  | macrophage_cDC_(inflam_fibroblast) |
| MYO7A |  |  | up | immune_CD8high_macrophagehigh_also_nonimmune |
| MOV10 |  |  | up | broad |
| HIRA |  | down |  | broad |
| PGBD4 |  | down |  | broad |
| ADSL | down |  |  | broad |
| GBE1 |  | down |  | malignant_enriched_noTcell |
| ARRDC4 | up |  |  | broad_noTcell |
| SLC45A4 |  | down |  | no_immune |
| FANK1 |  | up |  | Treg_malignant |
| LAMP5 |  |  | down | pDC_fibroblast |
| SLC2A10 |  |  | down | fibroblast_enriched_no_immune |
| MMRN1 |  | up |  | endothelial_no_immune |
| TFAP2A | down |  | up | malignant_enriched_no_immune |
| NKD1 | up |  |  | CRC_fibroblast_enriched_no_immune |
| S100A13 | down |  |  | no_immune |
